# Aggregation-related quenching of LHCII in liposomes revealed by single-molecule spectroscopy

**DOI:** 10.1101/2020.12.06.413419

**Authors:** Marijonas Tutkus, Jevgenij Chmeliov, Gediminas Trinkunas, Parveen Akhtar, Petar H. Lambrev, Leonas Valkunas

## Abstract

Incorporation of membrane proteins into reconstituted lipid membranes is a common approach for studying their structure and function relationship in a native-like environment. In this work, we investigated fluorescence properties of liposome-reconstituted LHCII. By utilizing liposome labelling with the fluorescent dye molecules and single-molecule microscopy techniques, we were able to study truly liposome-reconstituted LHCII and compare them with bulk measurements and liposome-free LHCII aggregates on bound surface. Our results showed that fluorescence lifetime in bulk and of that for single liposome measurements were correlated. The fluorescence lifetimes of LHCII were shorter for liposome-free LHCII than for reconstituted LHCII. In the case of liposome-reconstituted LHCII, fluorescence lifetime showed dependence on the protein density reminiscent to concentration quenching. The dependence of fluorescence lifetime of LHCII on the liposome size was not significant. Our results demonstrated that fluorescence quenching can be induced by LHCII-LHCII interactions in reconstituted membranes, most likely occurring via the same mechanism as photoprotective non-photochemical quenching in vivo.

## Introduction

Oxygenic photosynthetic organisms encounter large variations in the light intensity, which can change abruptly between extreme values. Evidently, light-harvesting needs to be finely regulated to achieve optimal efficiency of solar energy capture under low light conditions yet protect the organism from photo-damage under excess light. Indeed, under high light conditions, when reaction centres are progressively saturated, the excess absorbed energy might lead to the formation of potentially harmful singlet oxygen species. To avoid this damage, a self-protection mechanism using heat dissipation, called non-photochemical fluorescence quenching (NPQ), is evolved [1].

In plants, NPQ primarily occurs in the major light-harvesting complex II (LHCII). This process involves specific conformational changes of LHCII as well as some variations in the supramolecular organization of these proteins within the thylakoid membranes [2–4]. Similar fluorescence quenching could be triggered *in vitro* by removing detergent from the samples with solubilized LHCII complexes, thus inducing their aggregation [5]. Recent studies indicated that similar aggregation-related quenching mechanism is also evident in the native photosynthetic membrane [6,7]. It is thought that *in vitro* aggregation causes quenching via chlorophyll–chlorophyll contacts that are less feasible when LHCII is embedded in the lipid membrane [8]. In such aggregates, some light-harvesting complexes might switch into different conformational states responsible for the fast excitation energy quenching, as indicated on a single-LHCII level using single-molecule (SM) spectroscopy [9,10]. The presence of such trapping centers within the aggregate significantly reduces the observed excitation lifetime [6].

One of the widely used approaches to study properties of the membrane proteins in a native-like environment is their incorporation into liposomes [11]. Recently we showed, however, that application of this method to the LHCII was not as straightforward as it was assumed before. During the protein reconstitution, not only proteoliposomes were formed, but empty liposomes as well as non-reconstituted aggregated LHCII complexes were also present [12]. The resulting mixture made simple bulk measurements not very informative, and application of the SM microscopy method helped us to distinguish the fluorescence signals of LHCII originating in the proteoliposomes from that of the free LHCII aggregates.

In this work, we further employed surface immobilization of LHCII proteoliposomes and supplemented it with the time-resolved confocal fluorescence microscopy to study fluorescence decay kinetics in each distinct LHCII proteoliposome. In particular, we examined LHCII complexes reconstituted into liposomes at varying lipid:protein (L:P) ratios. Proteoliposomes exhibited a high degree of heterogeneity with respect to their size, protein density (PD) and fluorescence yield, even after separation into fractions by density gradient ultracentrifugation. Results of these experiments clearly demonstrated that the fluorescence yield and the fluorescence lifetime are gradually reduced with increasing protein density to values comparable with those of strongly quenched aggregates in the detergent-free solutions. This indicated the formation of the variously sized LHCII aggregates with some trapping centers upon reconstitution into liposomes and further supported the idea that similar aggregation-related quenching governs natural NPQ in the native membrane environment.

## Materials and methods

### Sample preparation

LHCII proteoliposome sample preparation was the same as in our previous work [12,13]. LHCII complexes were isolated from pea (Pisum sativum) plants, frozen in liquid N_2_ and stored at −80 °C until reconstitution. Liposomes were prepared from plant thylakoid lipids, the lipophilic fluorescent dye DiI (Invitrogen), the biotinylated lipid (1,2-distearoyl-sn-glycero-3-phosphoethanolamine-N-[biotinyl(poly- (ethylene glycol))-2000]) (Avanti Polar Lipids). Isolated LHCII complexes were reconstituted into liposomes using detergent destabilization. The trimeric LHCII complexes dissolved in β-DDM were added drop-wise to a suspension of liposomes at concentration of 5 mg/mL, while agitating continuously, to obtain a mixture of molar L:P of 500:1 and 1500:1. The L:P ratio of this reconstitution mixture was estimated taking into account that LHCII trimer contains 14 Chls per monomer. The detergent was then removed by repeated incubation of the sample with absorbent beads. Proteoliposome fractions of different densities and L:P were separated by density Ficoll PM-400 (GE Healthcare) gradient ultracentrifugation. Four to five colored band fractions of different densities were collected.

Silanized and PEGylated (methoxy-PEG and biotin−PEG) glass slides (#1.5, Menzel Glaser) were prepared in the same way as described previously [14]. These glass slides were assembled into the six-channel flow cells (sticky-Slide VI 0.4, IBIDI, Germany). Next channel was filled with the reconstitution buffer (300 μL) and incubated with neutravidin (nAv, A-26666, Molecular Probes) solution (0.02 mg/mL) for at least 3 min. Free nAv was removed by washing the channel with the reconstitution buffer (300 μL). Next, proteoliposomes were injected into the channel at low concentration and incubated until sufficient liposome density was achieved. Free proteoliposomes were removed by washing the channel with the reconstitution buffer (300 μL). Therefore, there was low probability to have detergent in the solution during the microscopy experiment. After this step, the sample was ready for microscopy measurements.

### Microscopy measurements and data analysis

A home-build stage-scanning confocal fluorescence microscope was used to acquire fluorescence images and fluorescence decays at room temperature. The sample was excited at 635 nm utilizing a picosecond diode laser (BHL-600, Becker & Hickl) with a pulse width of 60 ps and a repetition rate of 50 MHz, ∼0.05 µW (∼10 W/cm^2^), and at 532 nm (Crystalaser) with ∼0.2 µW (∼140 W/cm^2^). Laser powers were estimated at the plane of the sample. The fluorescence emission was collected with the high-NA oil-immersion objective (100×, 1.4 NA, Nikon), passed through a four-band emission filter (446/510/ 581/703, Semrock), and then directed to the dichroic mirror where the light was split into green and red components. The intensity of the green component was measured with one avalanche photodiode (APD) (tau-SPAD, PicoQuant, Germany), and the intensity of the red component was measured with a second identical APD. Intensity counts from the APDs were read out by an NI card (SCB-68, NI) connected to the FPGA card (PCIe-7851R, NI).

The wavelength-integrated fluorescence was measured with the same red-channel APD as for intensity measurements. A time-correlated single photon counting (TCSPC) device (Becker & Hickl Simple-Tau 140) was connected to the APD and allowed us to acquire triggered by pulsed laser excitation arrival time of the detected photons. Integration time for fluorescence decay collection was 2 s. The dichroic mirror and the fluorescence emission filtered out most of the excitation light.

Sample was scanned with the piezo stage (P-733.2CD, PI, Germany) with the controller (E-712.3CDA, PI, Germany). Control of the scanning and readout was achieved by custom-written LabVIEW (NI) software. Pixel dwell time was set to 5 ms, and size of the image was set to 200 × 200 pixels.

Image analysis was performed using custom-written software (Igor Pro 6.37) as described previously [15]. To extract the integrated fluorescence intensity, the intensity of the detected spots was fitted with a two-dimensional symmetric Gaussian function. Co-localization distance was set to 3 pixels. The PD parameter was expressed as the ratio of fluorescence intensity of LHCII and that of liposome dyes.

The fluorescence lifetimes were obtained by an exponential re-convolution fit using an instrument response function (IRF) measured at the excitation wavelength of 635 nm. The full width at half maximum (FWHM) of the IRF was ∼400 ps and was dominated by the timing error of the detector. The quality of the fitting procedure was evaluated from the lack of structure in the fit residuals and their auto-correlation function.

To convert liposome intensities into diameter in nanometers, we followed a previously described procedure [16]. We measured average liposome sizes of the control liposome sample without protein using a DLS machine (Zetasizer μV, Malvern Analytical). Our control liposomes had an average diameter of 106 nm. On our microscope setup, we imaged the same control liposomes as in DLS, obtained integrated intensities, took the square root of the intensities, and calculated the average of these distributions.

## Results and discussion

LHCII proteoliposomes have been considered as a good model system of thylakoids. This system allows one to study LHCII structural organization in a native-like lipid environment while eliminating spectral and temporal overlap of many processes occurring *in vivo* (e.g. the presence of spectrally distinct photosystems I and II as well as, under high light conditions, centers of both photochemical and non-photochemical excitation trapping) [17]. However, we demonstrated that this system is still not ideal in terms of the heterogeneity because a substantial fraction of LHCII in such samples remains liposome-free [12], meaning that the ordinary bulk spectroscopic measurements might represent the signal coming from both the proteoliposomes and the free (non-reconstituted) LHCII oligomers. Nonetheless, application of the SM microscopy techniques to LHCII studies in the dye-labelled liposomes allowed co-localization-based separation of liposome-free and liposome-reconstituted LHCII [12,18]. Therefore, in this work we step up with employing labeled LHCII proteoliposomes to model a natural light-harvesting system by taking into account fluorescence lifetime characteristics for the LHCII complexes.

We prepared two LHCII proteoliposome samples with different L:P ratios of the mixture, namely 500:1 and 1500:1. These samples were purified using the sucrose gradient centrifugation to remove the major part of the non-reconstituted LHCII. After this step, we obtained gradient bands with different mean L:P determined for each sample (Fig. 1A). From the 500:1 sample, we obtained gradient bands with L:P ratio of 30:1 (B4) and 40:1 (B3), while the bands obtained from the 1500:1 sample exhibited the following L:P ratios—250:1 (B3), 70:1 (B4), and 67:1 (B5). In the text below, we focused mainly on the results from the B3–B5 bands of the 1500:1 sample because they showed more differences than those of the 500:1 sample. For comparison, the results of the 500:1 sample bands are provided in the Supporting Information (SI).

**Figure 1.**
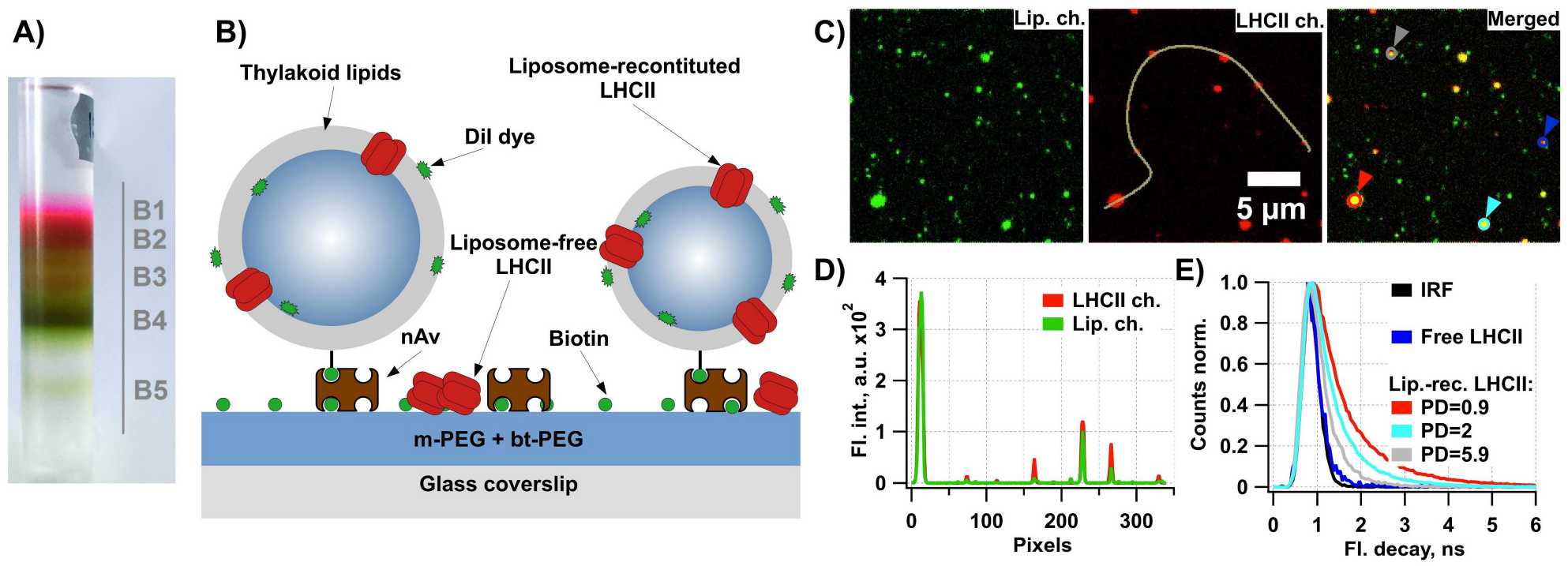
The assay of single immobilized LHCII proteoliposomes and representative results. **A)** Photograph of the test tube after gradient centrifugation with indicated 5 different gradient bands (B1 – B5). **B)** Illustration of the immobilization of the proteoliposomes containing biotinylated lipids on the m-PEG/bt-PEG (10:1) modified surface via neutravidin (nAv). Liposomes were labeled using DiI dye during their preparation. **C)** Representative confocal fluorescence microscopy images acquired at 532 and 635 nm excitation (liposome dye and LHCII channels, respectively) and merged. **D)** Graph showing line-scans (along the path indicated on the LHCII ch. image in the panel **C** taken over several liposomes in the LHCII and liposome channels. **E)** Graph showing IRF and illustrative fluorescence decay kinetics of liposome-free and reconstituted LHCII of three different protein densities (PD), taken from the spots indicated with colored triangles in the merged channel image of panel **C**. For interpretation of the references to color in this figure legend, the reader is referred to the web version of this article.

The immobilization and labeling of LHCII proteoliposomes were performed using the same techniques as in our previous work [12]. Briefly, LHCII proteoliposomes, prepared from the mixture of the thylakoid lipids, biotinylated lipids and DiI membrane dye, were immobilized on the surface of silanized and PEGylated (10% bt-PEG) glass coverslip via nAv (Fig. 1B). The coverslip was mounted on the bottom flowcell’s surface before the immobilization. We performed confocal fluoresce lifetime imaging microcopy on the five aforementioned samples obtained from the gradient bands. Strong differences in the absorption spectra of the LHCIIs and the dye molecules provided us with the opportunity to selectively excite either LHCIIs (both reconstituted into the liposomes and the free trimers and/or aggregates remaining in the sample) or liposomes (both containing LHCII and empty). The obtained images of the LHCII channel (at 635 nm excitation) and liposome channel (at 532 nm excitation) are shown in Fig. 1C. By comparing these images and overlapping them to perform co-localization analysis, we were able to discriminate between the fluorescing spots representing the empty liposomes, free LHCIIs, and liposome-reconstituted LHCIIs. As reported previously, all of the examined samples showed significant fractions of the non-reconstituted LHCII. The percentage of liposome-free LHCII was dependent on L:P ratio of the gradient band—bands with lower L:P ratio exhibited more liposome-free LHCII (SI Table 1).

Integrated liposome fluorescent spot intensity is proportional to the surface area of the liposome. Thus it can be converted into diameter, expressed in nanometers, using a specific conversion factor [16]. This factor was obtained by comparing DLS-measured liposomes sizes and confocal microscope-measured square-rooted liposomes intensities of the control liposome sample without LHCII (SI Fig. 1). Control liposomes in DLS measurements showed a mean diameter of ∼ 106 nm. As reported previously, visual inspection and intensity analysis of the acquired images indicated up-concentration of LHCII in smaller liposomes (Fig. 1D).

To ensure that the surface immobilization did not introduce any artifacts in fluorescence lifetime kinetics of LHCII, we performed control measurements with the detergent-solubilized LHCII trimers, immobilized on the PLL covered glass coverslip surface (SI Fig. 2A). We acquired their confocal fluorescence images (SI Fig. 2B), detected fluorescent LHCII spots in them, and performed registration of fluorescence lifetime decays from each of the detected LHCII spots. Results of these measurements showed that the fluorescence decays of the surface-immobilized LHCII can be fitted using a single-exponential function, and their mean fluorescence lifetime was ∼3 ns (SI Fig. 2C). The obtained mean value of the lifetime was similar to the mean fluorescence lifetime in an ensemble of solubilized complexes [19] and to that of the single immobilized LHCII subject to the <50 W/cm^2^ excitation power, when any non-linear annihilation effects are absent [20].

**Figure 2.**
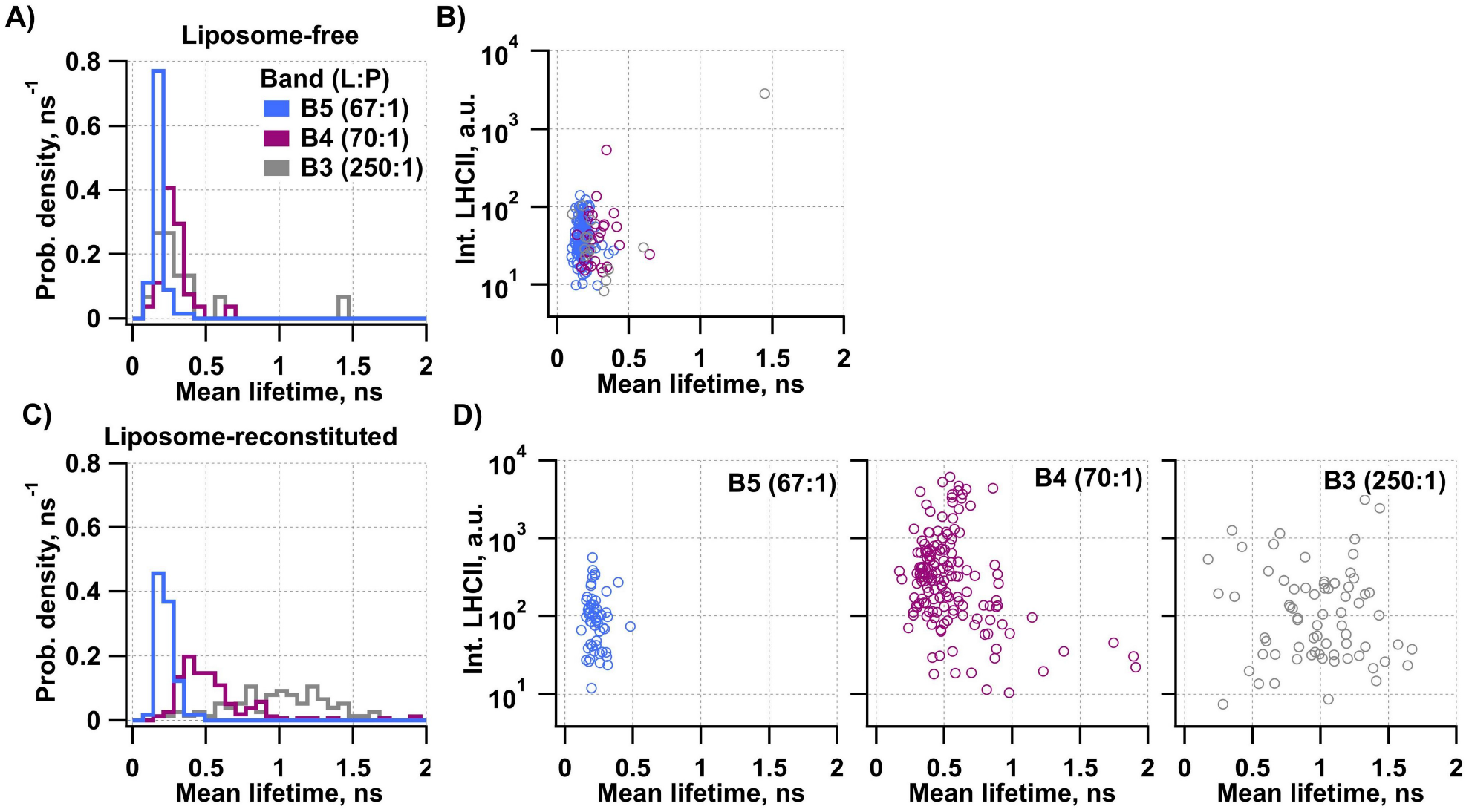
Results of fluorescence lifetime imaging microcopy. Graphs showing distribution of fluorescence lifetimes obtained by the single exponential fitting of **A)** liposome-free and **C)** reconstituted LHCII of three samples with different lipid to protein (L:P) ratios. Panels **B** and **D** demonstrate correlation of fluorescence lifetimes of free and liposome-reconstituted LHCII of three samples with different L:P. For interpretation of the references to color in this figure legend, the reader is referred to the web version of this article.

During the LHCII proteoliposome measurements, we collected fluorescence decay kinetics from each of the detected LHCII spots regardless of whether it was liposome-free or reconstituted (Fig. 1E). Co-localization analysis of the lipid-dye and LHCII fluorescence allowed us to distinguish between these two cases. The absolute majority of the liposome-free LHCIIs exhibited very fast excitation decay kinetics with the mean lifetimes about 200 ps (see blue kinetics in Fig. 1E and SI Table 2), which was similar to those obtained in the LHCII aggregates formed by removing detergent [5,6,19,21]. This observation suggested that the majority of the liposome-free LHCII in our samples was in the form of the relatively large LHCII oligomers. On the other hand, fluorescence kinetics from the liposome-reconstituted LHCII exhibited various decay rates (Fig. 1E).

To characterize individual proteoliposomes, we used several numerical quantities. First, we calculated the ratio of the integrated fluorescence intensities in the LHCII and dye channels to estimate the mean protein density (PD) of LHCII in each liposome. Next, fluorescence decay kinetics for each detected LHCII spot was fitted with the convolution of the IRF and a single-exponential function. Fitting with double-exponential function gave very similar results and did not reduce the residuals of fitting significantly, indicating that the faster decay components, if present, were far beyond the time resolution of our measurements.

The detected mean excitation lifetimes, obtained for different liposome-free LHCII oligomers in different samples, are presented in SI Fig. 3A, and their statistical distribution is shown in Fig. 2A and SI Fig 4A. We can see that in most oligomers, the lifetimes ranged from 0.1 to 0.3 ns, although several spots exhibited longer fluorescence decay kinetics. This variation of the average lifetimes between different samples indicated that LHCII aggregates of various sizes and possibly different amount of the generated NPQ traps were formed. Meanwhile, results of liposome-reconstituted LHCII fluorescence decays analysis showed that only in the B5 gradient band the majority LHCII proteolipomes exhibited similarly short fluorescence lifetime of about 0.23 ns (see Fig. 2C, SI Fig. 3C and SI Table 2). In contrast, proteoliposomes from the B4 and B3 gradient bands showed significantly slower average fluorescence lifetimes of ∼ 0.56 ns and 1.01 ns, respectively. Similar effect was also demonstrated by the bulk measurements [13], though in our work elimination of the liposome-free LHCIIs from the analysis resulted in the slightly longer mean lifetimes.

**Figure 3.**
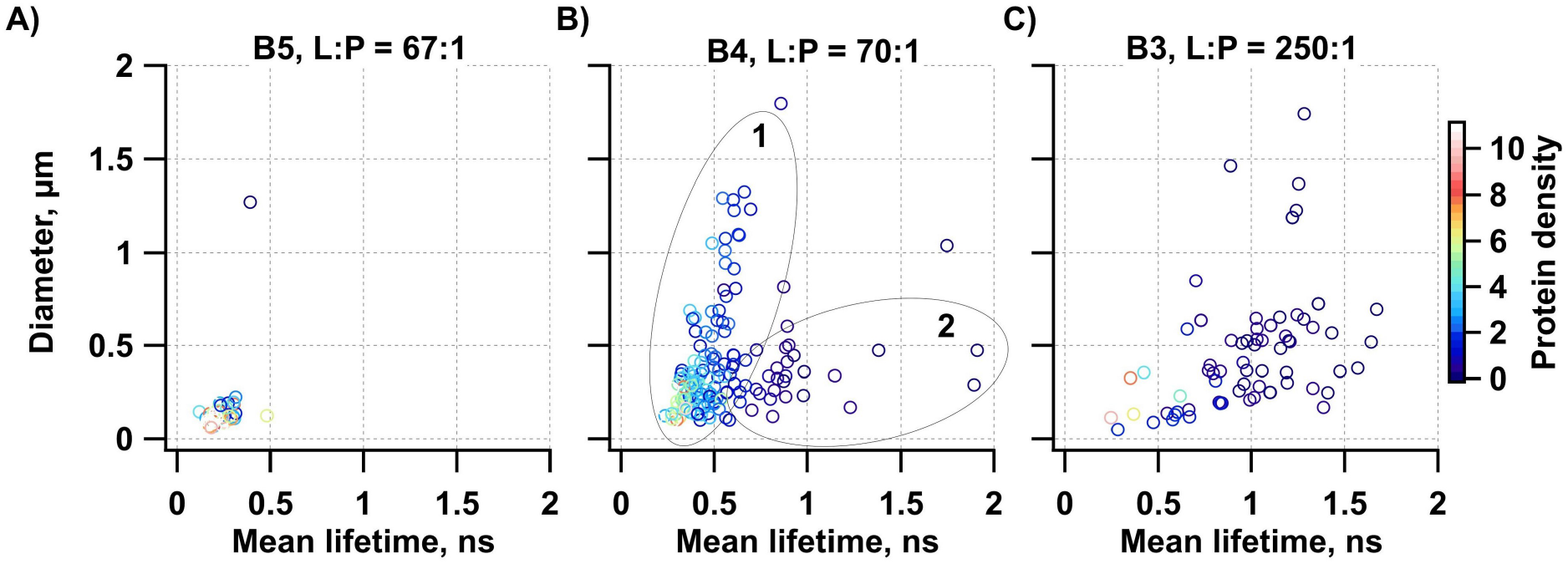
Relation between the fluorescence lifetime, liposome diameter, and protein density of liposome-reconstituted LHCII from three LHCII proteoliposome samples with different lipid to protein ratios (L:P): **A)** 67:1 (B5), **B)** 70:1 (B4), and **C)** 250:1 (B3). These three samples were obtained from gradient bands of 1500:1 proteoliposome sample. Color-code represents the protein density. For interpretation of the references to color in this figure legend, the reader is referred to the web version of this article.

**Figure 4.**
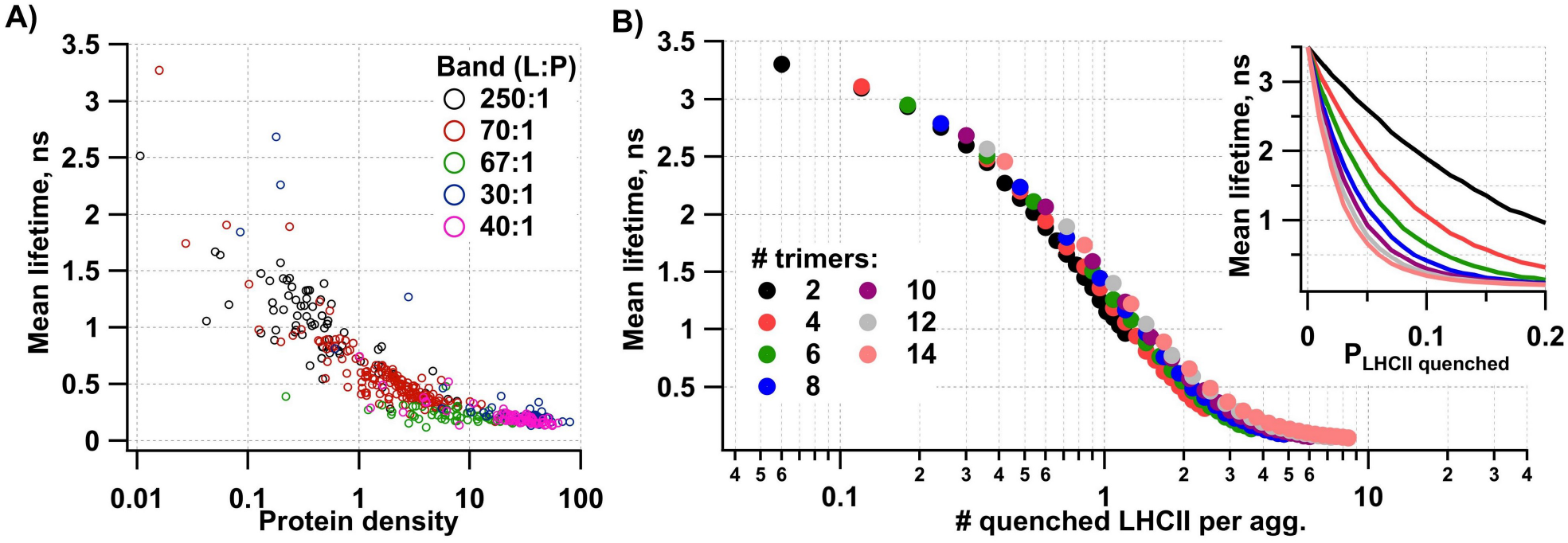
Fluorescence mean lifetime dependence of liposome-reconstituted LHCII on protein density. **A)** Graph showing dependence of the mean fluorescence lifetime of liposome-reconstituted LHCII on the protein density of the proteoliposome for different gradient band samples. **B)** Graphs showing results of LHCII aggregation modeling in bulk. The inset of this panel shows the dependence between the mean excitation lifetime in the aggregates containing 2–14 LHCII trimers and the probability for any complex in the aggregate to be in the quenched state, P_LHCII quenched_ [6,27]. All distinct curves overlap remarkably when plotted against the mean number of the quenched complexes per aggregate (main graph), which is defined as the aggregate size times the probability for the complex to be in the quenching state. For interpretation of the references to color in this figure legend, the reader is referred to the web version of this article.

In all the samples, the detected LHCII fluorescence intensities have exhibited a rather significant variation (see SI Fig. 3B and D). However, they did not show any clear correlation with the mean fluorescence lifetime (see Fig. 2B and 2D for free and reconstituted LHCII, respectively). This could also be explained by an existence of variously sized LHCII aggregates and their great abundance in proteoliposomes. Also, this result suggested that not all LHCIIs in the liposome form a single cluster. Indeed, the fact that there were a number of proteoliposomes with the same mean lifetime and very broad—covering several orders of magnitudes—distribution of the fluorescence intensities suggested that in those liposomes various number of similar LHCII oligomers were formed. In this case, different clusters would sum-up, resulting in different intensity values for different liposomes, but their fluorescence lifetimes would remain independent.

To figure out what effect has the PD (SI Fig. 3E) and liposome size (SI Fig. 3F) on the LHCII fluorescence lifetime, we plotted the liposome diameter versus the fluorescence lifetime over color-code represented PD (Fig. 3 and SI Fig. 5). For the B5 gradient band, all of these three parameters demonstrated the smallest variations (Fig. 3A). In this band, all the liposomes were small (< 0.3 µm in diameter), had high-to-moderate PD, and LHCII therein were highly quenched (τ < 0.3 ns). The same plot for the B4 gradient band showed wide distribution of liposome sizes, PD, and LHCII fluorescence lifetimes (Fig. 3B). This funnel-shaped plot revealed two populations of LHCII proteoliposomes in this gradient band: one with high PD and broad distribution of liposome sizes (vertical, population 1), and another with low PD and smaller liposome sizes (horizontal, population 2). The LHCIIs in the first population were more quenched than in the second population, and also showed broader distribution of the fluorescence lifetimes. The plot for the B3 gradient band had the lowest PD and the least quenched LHCIIs (Fig. 3C).

Based on these results we could conclude that the level of LHCII aggregation, reflected by the mean fluorescence lifetime, is determined by the PD of LHCII proteoliposomes rather than liposome size. In addition to that, our results showed that the gradient centrifugation could relatively well sort out the proteoliposomes with highly aggregated LHCII (B5) from less aggregated LHCII (B3). However, in the B4 band we found both types of LHCII aggregates. The vague separation could be explained by the fact that the gradient centrifugation separates proteoliposomes according to their total mass, which is determined by the liposome mass and the mean protein density therein. Therefore, small liposomes with high PD and big liposomes with lower PD are collated in the same gradient band. As mentioned earlier, the gradient centrifugation procedure did not allow us fully eliminate liposome-free LHCII. Therefore, it was crucial to employ labeling of the liposome membrane with a fluorescent dye, which was utilized for co-localization-based sorting in the data analysis for liposome-free and reconstituted LHCIIs.

The results presented above clearly showed the high level of heterogeneity in studied proteoliposomes samples, even after gradient centrifugation: the liposomes themselves varied in size, and even the liposomes of the same size had various amount of the incorporated LHCII complexes. In the latter case, higher PD generally corresponded to the faster excitation decay kinetics (*cf*. color variations of the dots along the horizontal cuts in Fig. 3). To further investigate this behavior, we plotted the fluorescence lifetime versus PD for all the studied samples (see Fig. 4A). Interestingly, despite the high degree of heterogeneity of the measured properties between the gradient bands and within each band, upon presenting the data from all the proteoliposomes on the same graph, all data points laid along a continuous sigmoidal curve. At a low PD, only several LHCII trimers per liposome are expected, therefore the curve in Fig. 4A extrapolated to the mean lifetimes of ∼3 ns, similar to that observed in LHCII trimers. On the other hand, at high PD the reconstituted LHCII trimers form large dense clusters and the *τ*(PD) curve converged at ∼200 ps. This value is limited by the FWHM of the IRF of our setup, thus any faster decay kinetics were not resolved.

This dependency resembled the old-standing chlorophyll concentration quenching curve [22,23]. One difference, however, is that for the liposomes we observe variation of PD over 4 orders of magnitude, while for chlorophylls this dependence lasted over 2 orders of magnitude of the chlorophyll concentration. We would like to note, however, that PD (i.e. the ratio of LHCII and lipid-dye fluorescence intensities) represents only some effective, or average, protein concentration in the liposome: the fluorescence intensity of LHCII depend on their clustering, therefore, higher fluorescence intensity would be measured when LHCII were well-separated within the liposome, compared to the case when all of them formed a cluster. Indeed, the calculated PD would be different for these two cases, although there would be the same number of LHCII complexes per liposome. Such uncertainty might be the reason for apparent much wider variations in PD compared to the actual LHCII concentration (i.e. amount of LHCII complexes per liposome surface area).

Aggregation-related excitation quenching of light-harvesting complexes results from their intrinsic ability to switch between several distinct conformational states [24]. As revealed by means of the SM spectroscopy, each single LHCII trimer undergoes reversible transitions between the fluorescent and dark (or quenching) states [10,25,26]. Therefore, in a large aggregate there is always a higher probability that some of the constituting complexes will be in the quenched conformation, thus excitation energy migration through the aggregates to such excitation traps will result in the strongly quenched fluorescence decay kinetics [6,27,28].

In the inset of Fig. 4B we show how the mean excitation lifetime in variously sized aggregates drops down with the increase of probability for any complex in the aggregate to be in the quenched state (coarse-grained calculations were performed assuming inter-complex energy transfer rate of (25 ps)^−1^ and the quenching time by the traps of 50 ps, see [6,27] for more details). All these distinct curves, obtained for various aggregates, however, overlapped remarkably when plotted against the mean number of the quenched complexes per aggregate (Fig. 4B). The latter parameter is defined as the aggregate size times the probability for the complex to be in the quenching state (P_LHCII quenched_). The sigmoidal shape of the obtained dependence also resembled the one obtained for the *τ*(PD) curve describing the mean excitation lifetime in the liposome. This fact again emphasized the aggregation-related nature of the observed fluorescence lifetime variations in the proteoliposomes.

## Conclusion

Previously performed bulk fluorescence spectroscopy measurements of LHCII proteoliposomes at 77 K temperature showed short fluorescence lifetimes and an increased shoulder around 700 nm in the fluorescence spectrum that indicated the presence of LHCII aggregates [13]. This aggregation degree was dependent on the L:P ratio of the sample. However, in that bulk assay it was not clear whether the observed effects are due to remaining liposome-free LHCII oligomers or liposome-reconstituted LHCII. In this work, we aimed to better understand processes that occur in the LHCII in proteoliposomes. We utilized co-localization between the membrane dye and the LHCII to discriminate between liposome-free and reconstituted LHCIIs. By employing our previously developed and characterized assay, we observed fluorescence quenching in LHCII in natural-like environment. Our results showed that the fluorescence lifetime in bulk and single-liposome measurements were correlated: samples with lower L:P ratio on average had higher PD and also shorter average fluorescence lifetime. The fluorescence lifetimes of LHCII were shorter for liposome-free LHCII than for liposome-reconstituted LHCII. For all the samples we obtained a unique dependence between the fluorescence lifetime and the mean PD of the liposomes, defined in this work as the ratio of the LHCII and lipid dyes fluorescence intensities. Despite exhibiting high level of heterogeneity in almost all observable parameters – the liposome size, fluorescence intensity of LHCII and liposome dye, and mean fluorescence lifetime of LHCII – all the obtained results confirmed the aggregation-related quenching of LHCII proteins in protein-dense liposomes.

Moreover, our single-proteoliposome results showed that the quenching level and thus the size of the LHCII aggregate was dependent on lipid-to-protein ratio of the LHCII proteoliposome sample. Generally, it is difficult to prepare liposomes without LHCII aggregates [29]. However, gradient centrifugation might help to sort out proteoliposomes with different degree of LHCII aggregation. Previous reports demonstrated that LHCII were unquenched in the nanodiscs [30]. However, in the nanodiscs LHCII were constrained to single protein copy per nanodisc. Recent report claims that it was possible to reach dark-adapted-like LHCII in proteoliposomes as well [31,32]. However, it is not clear what is the degree of liposome-free LHCII in those samples, and our co-localization-based microscopy assay might be applied for this purpose. Our observed tendency of aggregation in lipid bilayer might reflect the situation in thylakoid membranes where LHCII is free to diffuse laterally.

## Acknowledgment

This work was supported by the Lithuanian Research Council (no. S-LZ-19-3) (MT, JC, and LV) and World Federation of Scientist national scholarship awarded for MT. P.H.L. acknowledges support from the National Research, Development and Innovation Fund (NN-124904). We also thank Mikas Vengris, Danielis Rutkauskas for their excellent technical assistance and Fanni Görföl for help with sample preparation.

## References

[1] B. Demmig-Adams, G. Garab, W. Adams III, Govindjee, eds., Non-Photochemical Quenching and Energy Dissipation in Plants, Algae and Cyanobacteria, Springer Netherlands, Dordrecht, 2014. https://doi.org/10.1007/978-94-017-9032-1.

[2] C.D.P. Duffy, L. Valkunas, A. V. Ruban, Light-harvesting processes in the dynamic photosynthetic antenna, Phys. Chem. Chem. Phys. 15 (2013) 18752–18770. https://doi.org/10.1039/c3cp51878g.

[3] E. Belgio, E. Kapitonova, J. Chmeliov, C.D.P. Duffy, P. Ungerer, L. Valkunas, A. V. Ruban, Economic photoprotection in photosystem II that retains a complete light-harvesting system with slow energy traps, Nat. Commun. 5 (2014) 4433. https://doi.org/10.1038/ncomms5433.

[4] A. V. Ruban, Nonphotochemical Chlorophyll Fluorescence Quenching: Mechanism and Effectiveness in Protecting Plants from Photodamage, Plant Physiol. 170 (2016) 1903–1916. https://doi.org/10.1104/pp.15.01935.

[5] P. Horton, A. V. Ruban, D. Rees, A.A. Pascal, G. Noctor, A.J. Young, Control of the light-harvesting function of chloroplast membranes by aggregation of the LHCII chlorophyll-protein complex, FEBS Lett. 292 (1991) 1–4. https://doi.org/10.1016/0014-5793(91)80819-O.

[6] J. Chmeliov, A. Gelzinis, E. Songaila, R. Augulis, C.D.P. Duffy, A. V. Ruban, L. Valkunas, The nature of self-regulation in photosynthetic light-harvesting antenna, Nat. Plants 2 (2016) 16045. https://doi.org/10.1038/nplants.2016.45.

[7] J. Chmeliov, A. Gelzinis, M. Franckevičius, M. Tutkus, F. Saccon, A. V Ruban, L. Valkunas, Aggregation-Related Nonphotochemical Quenching in the Photosynthetic Membrane, J. Phys. Chem. Lett. 10 (2019) 7340–7346. https://doi.org/10.1021/acs.jpclett.9b03100.

[8] T. Barros, A. Royant, J. Standfuss, A. Dreuw, W. Kühlbrandt, Crystal structure of plant light-harvesting complex shows the active, energy-transmitting state, EMBO J. 28 (2009) 298–306. https://doi.org/10.1038/emboj.2008.276.

[9] T.P.J. Krüger, V.I. Novoderezhkin, C. Ilioaia, R. Van Grondelle, Fluorescence spectral dynamics of single LHCII trimers, Biophys. J. 98 (2010) 3093–3101. https://doi.org/10.1016/j.bpj.2010.03.028.

[10] M. Tutkus, J. Chmeliov, D. Rutkauskas, A. V. Ruban, L. Valkunas, Influence of the Carotenoid Composition on the Conformational Dynamics of Photosynthetic Light-Harvesting Complexes, J. Phys. Chem. Lett. 8 (2017) 5898–5906. https://doi.org/10.1021/acs.jpclett.7b02634.

[11] J.-L. Rigaud, D. Lévy, Reconstitution of Membrane Proteins into Liposomes, in: Methods Enzymol., Academic Press, 2003: pp. 65–86. https://doi.org/10.1016/S0076-6879(03)72004-7.

[12] M. Tutkus, P. Akhtar, J. Chmeliov, F. Görföl, G. Trinkunas, P.H. Lambrev, L. Valkunas, Fluorescence Microscopy of Single Liposomes with Incorporated Pigment-Proteins, Langmuir 34 (2018) 14410–14418. https://doi.org/10.1021/acs.langmuir.8b02307.

[13] P. Akhtar, F. Görföl, G. Garab, P.H. Lambrev, Dependence of chlorophyll fluorescence quenching on the lipid-to-protein ratio in reconstituted light-harvesting complex II membranes containing lipid labels, Chem. Phys. 522 (2019) 242–248. https://doi.org/10.1016/j.chemphys.2019.03.012.

[14] M. Tutkus, T. Rakickas, A. Kopūstas, Š. Ivanovaitė, O. Venckus, V. Navikas, M. Zaremba, E. Manakova, R. Valiokas, Fixed DNA Molecule Arrays for High-Throughput Single DNA– Protein Interaction Studies, Langmuir 35 (2019) 5921–5930. https://doi.org/10.1021/acs.langmuir.8b03424.

[15] M. Tutkus, T. Marciulionis, G. Sasnauskas, D. Rutkauskas, DNA-Endonuclease Complex Dynamics by Simultaneous FRET and Fluorophore Intensity in Evanescent Field, Biophys. J. 112 (2017) 850–858. https://doi.org/10.1016/j.bpj.2017.01.017.

[16] A.H. Kunding, M.W. Mortensen, S.M. Christensen, D. Stamou, A fluorescence-based technique to construct size distributions from single-object measurements: Application to the extrusion of lipid vesicles, Biophys. J. 95 (2008) 1176–1188. https://doi.org/10.1529/biophysj.108.128819.

[17] S. Farooq, J. Chmeliov, E. Wientjes, R. Koehorst, A. Bader, L. Valkunas, G. Trinkunas, H. van Amerongen, Dynamic feedback of the photosystem II reaction centre on photoprotection in plants, Nat. Plants 4 (2018) 225–231. https://doi.org/10.1038/s41477-018-0127-8.

[18] A.M. Hancock, S.A. Meredith, S.D. Connell, L.J.C. Jeuken, P.G. Adams, Proteoliposomes as energy transferring nanomaterials: enhancing the spectral range of light-harvesting proteins using lipid-linked chromophores, Nanoscale 11 (2019) 16284–16292. https://doi.org/10.1039/C9NR04653D.

[19] P. Akhtar, M. Dorogi, K. Pawlak, L. Kovács, A. Bóta, T. Kiss, G. Garab, P.H. Lambrev, Pigment interactions in light-harvesting complex II in different molecular environments, J. Biol. Chem. 290 (2015) 4877–4886. https://doi.org/10.1074/jbc.M114.607770.

[20] M.J. Gruber, J. Chmeliov, T.P.J. Krüger, L. Valkunas, R. van Grondelle, Singlet–triplet annihilation in single LHCII complexes, Phys. Chem. Chem. Phys. 17 (2015) 19844–19853. https://doi.org/10.1039/C5CP01806D.

[21] B. van Oort, A. van Hoek, A. V. Ruban, H. van Amerongen, Aggregation of Light-Harvesting Complex II leads to formation of efficient excitation energy traps in monomeric and trimeric complexes, FEBS Lett. 581 (2007) 3528–3532. https://doi.org/10.1016/j.febslet.2007.06.070.

[22] G.S. Beddard, G. Porter, Concentration quenching in chlorophyll, Nature 260 (1976) 366–367. https://doi.org/10.1038/260366a0.

[23] G.S. Beddard, S.E. Carlin, G. Porter, Concentration quenching of chlorophyll fluorescence in bilayer lipid vesicles and liposomes, Chem. Phys. Lett. 43 (1976) 27–32. https://doi.org/10.1016/0009-2614(76)80749-X.

[24] L. Valkunas, J. Chmeliov, T.P.J. Krüger, C. Ilioaia, R. Van Grondelle, How photosynthetic proteins switch, J. Phys. Chem. Lett. 3 (2012) 2779–2784. https://doi.org/10.1021/jz300983r.

[25] T.P.J. Krüger, C. Ilioaia, R. van Grondelle, Fluorescence Intermittency from the Main Plant Light-Harvesting Complex: Resolving Shifts between Intensity Levels, J. Phys. Chem. B 115 (2011) 5071–5082. https://doi.org/10.1021/jp201609c.

[26] G.S. Schlau-Cohen, H.Y. Yang, T.P.J. Krüger, P. Xu, M. Gwizdala, R. Van Grondelle, R. Croce, W.E. Moerner, Single-molecule identification of quenched and unquenched states of LHCII, J. Phys. Chem. Lett. 6 (2015) 860–867. https://doi.org/10.1021/acs.jpclett.5b00034.

[27] H. van Amerongen, J. Chmeliov, Instantaneous switching between different modes of non-photochemical quenching in plants. Consequences for increasing biomass production, Biochim. Biophys. Acta, Bioenerg. 1861 (2020) 148119. https://doi.org/10.1016/j.bbabio.2019.148119.

[28] P.H. Lambrev, F.J. Schmitt, S. Kussin, M. Schoengen, Z. Várkonyi, H.J. Eichler, G. Garab, G. Renger, Functional domain size in aggregates of light-harvesting complex II and thylakoid membranes, Biochim. Biophys. Acta, Bioenerg. 1807 (2011) 1022–1031. https://doi.org/10.1016/j.bbabio.2011.05.003.

[29] A. Natali, J.M. Gruber, L. Dietzel, M.C.A. Stuart, R. Van Grondelle, R. Croce, Light-harvesting Complexes (LHCs) cluster spontaneously in membrane environment leading to shortening of their excited state lifetimes, J. Biol. Chem. 291 (2016) 16730–16739. https://doi.org/10.1074/jbc.M116.730101.

[30] A. Pandit, N. Shirzad-Wasei, L.M. Wlodarczyk, H. Van Roon, E.J. Boekema, J.P. Dekker, W.J. De Grip, Assembly of the major light-harvesting complex II in lipid nanodiscs, Biophys. J. 101 (2011) 2507–2515. https://doi.org/10.1016/j.bpj.2011.09.055.

[31] E. Crisafi, M. Krishnan, A. Pandit, Time-resolved fluorescence analysis of LHCII in the presence of PsbS at neutral and low pH, BioRxiv (2018). https://doi.org/10.1101/456046.

[32] F. Azadi-Chegeni, M.E. Ward, G. Perin, D. Simionato, T. Morosinotto, M. Baldus, A. Pandit, Conformational dynamics of photosynthetic Light-Harvesting Complex II in a native membrane environment, BioRxiv (2018). https://doi.org/10.1101/288860.

