## Supplementary information for "Aggregation-related quenching of LHCII in liposomes revealed by single-molecule spectroscopy"

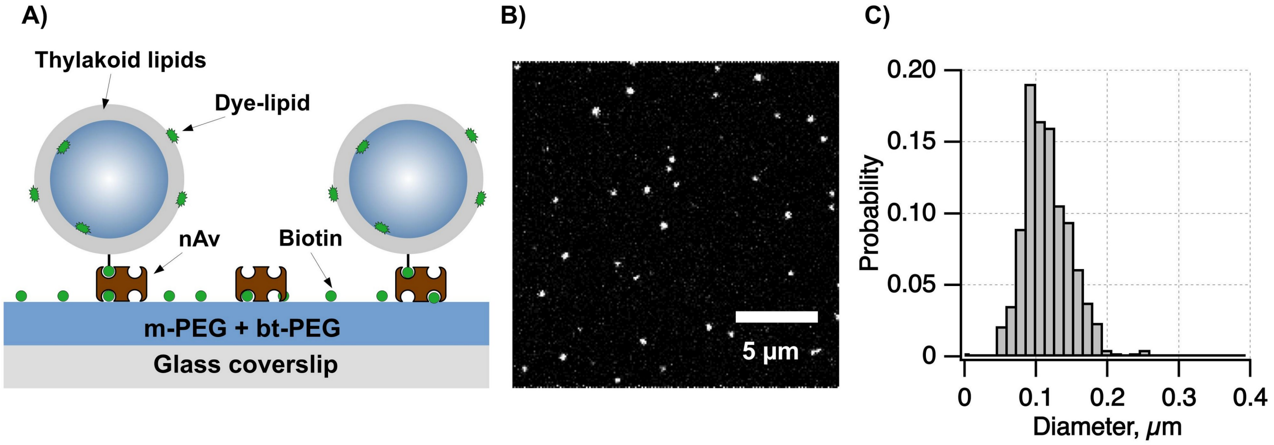


**SI Figure 1 | A)** Schematics illustrating the control liposome sample and immobilization strategy. **B)** Liposome channel confocal fluorescence image. **C)** Graph showing distribution of liposome sizes obtained from the confocal fluorescence images using 2D Gaussian fitting for more precise estimation of the total integrated intensity.

**SI Table 1** | Lipid to protein ratio (L:P), sample sizes and percentage of liposome-free LHCII in different LHCII proteoliposome samples.

| Band name | L:P mixture | L:P gradient band | Total number of detected LHCII spots | % of liposome-free LHCII |
|---|---|---|---|---|
| B3 | 1500:1 | 250:1 | 92 | 16.3 |
| B4 | 1500:1 | 70:1 | 184 | 14.7 |
| B5 | 1500:1 | 67:1 | 192 | 70 |
| B3 | 500:1 | 40:1 | 160 | 58.8 |
| B4 | 500:1 | 30:1 | 111 | 41.4 |

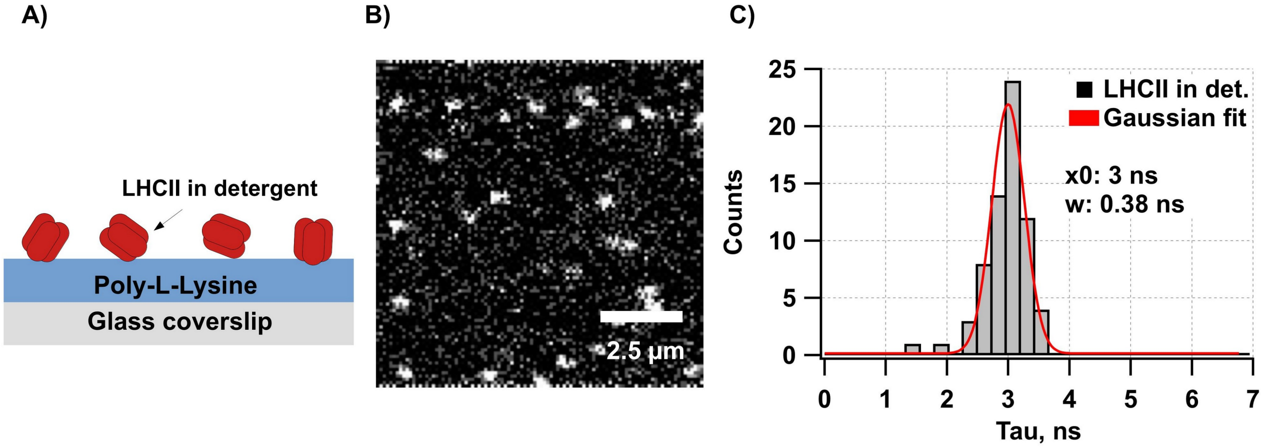


**SI Figure 2 | A)** Schematics illustrating LHCII in detergent micelles immobilized on the PLL modified glass coverslip surface. **B)** LHCII channel confocal fluorescence image of the immobilized LHCII trimers in detergent. **C)** Distribution of the fluorescence lifetimes (grey bars) with Gaussian fit (red curve). The Gaussian fitting results (center and width) are indicated at the figure legend.

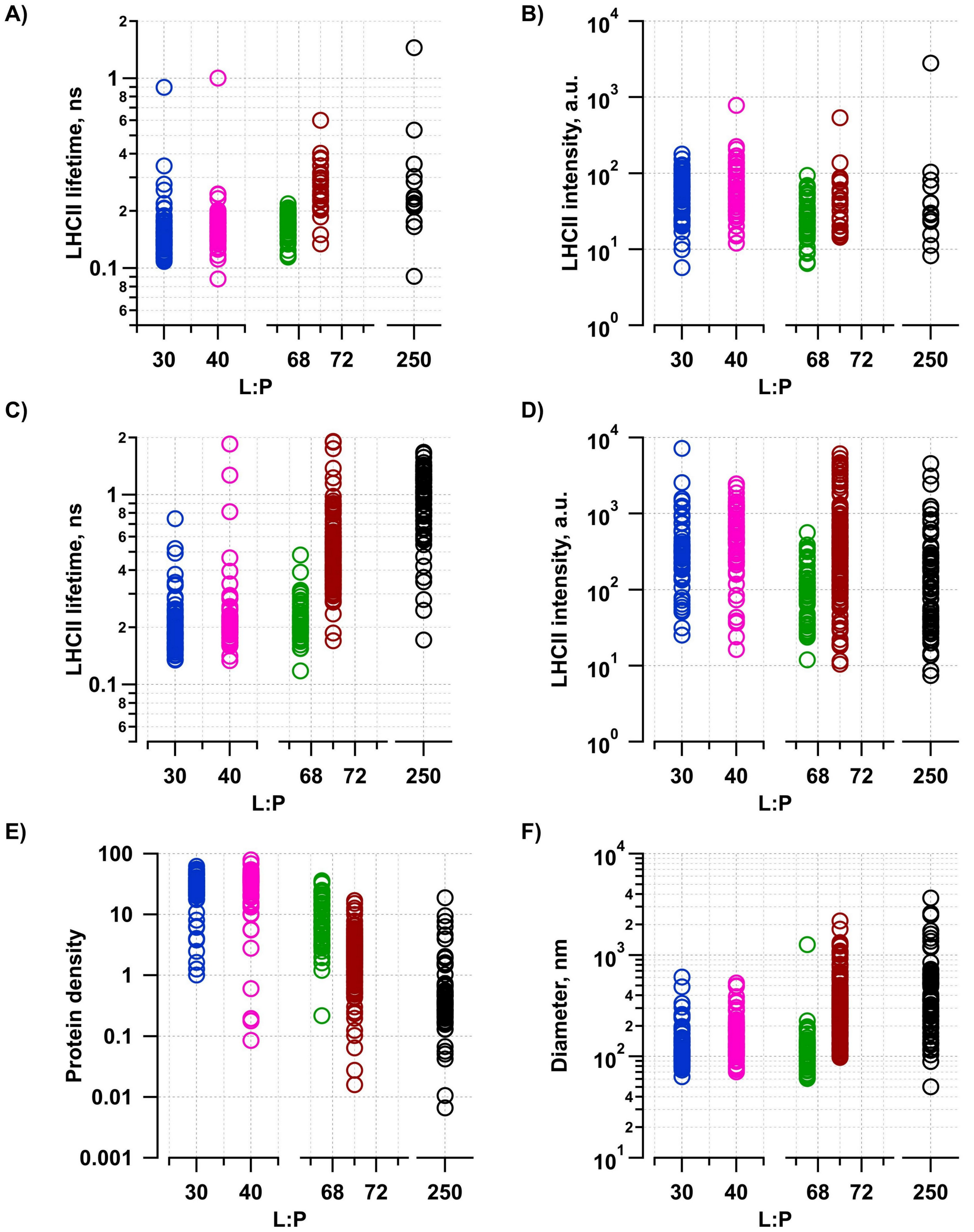


**SI Figure 3 |** Graphs showing results of the mean excitation lifetimes, obtained for **A)** liposome-free and **C)** -reconstituted LHCII in different lipid to protein (L:P) ratio samples. Graphs showing the detected LHCII fluorescence intensities for **B)** liposome-free and **D)** -reconstituted LHCII in different L:P ratio samples. Graphs showing results of **E)** protein density and **F)** liposome diameter obtained for liposome-reconstituted LHCII in different L:P ratio samples.

**SI Table 2** | Lipid to protein ratio (L:P), average and SD of fluorescence lifetime extracted using the single-exponential function fitting for different LHCII proteoliposome samples.

| Band name | L:P mixture | L:P gradient band | Liposome-reconstituted LHCII ⟨tau⟩ [ns] | Liposome-reconstituted LHCII SD tau [ns] | Liposome-free LHCII ⟨tau⟩ [ns] | Liposome-free LHCII SD tau [ns] |
|---|---|---|---|---|---|---|
| B3 | 1500:1 | 250:1 | 1.01 | 0.38 | 0.24 | 0.08 |
| B4 | 1500:1 | 70:1 | 0.56 | 0.35 | 0.29 | 0.1 |
| B5 | 1500:1 | 67:1 | 0.23 | 0.06 | 0.18 | 0.04 |
| B3 | 500:1 | 40:1 | 0.22 | 0.1 | 0.17 | 0.11 |
| B4 | 500:1 | 30:1 | 0.34 | 0.46 | 0.19 | 0.13 |

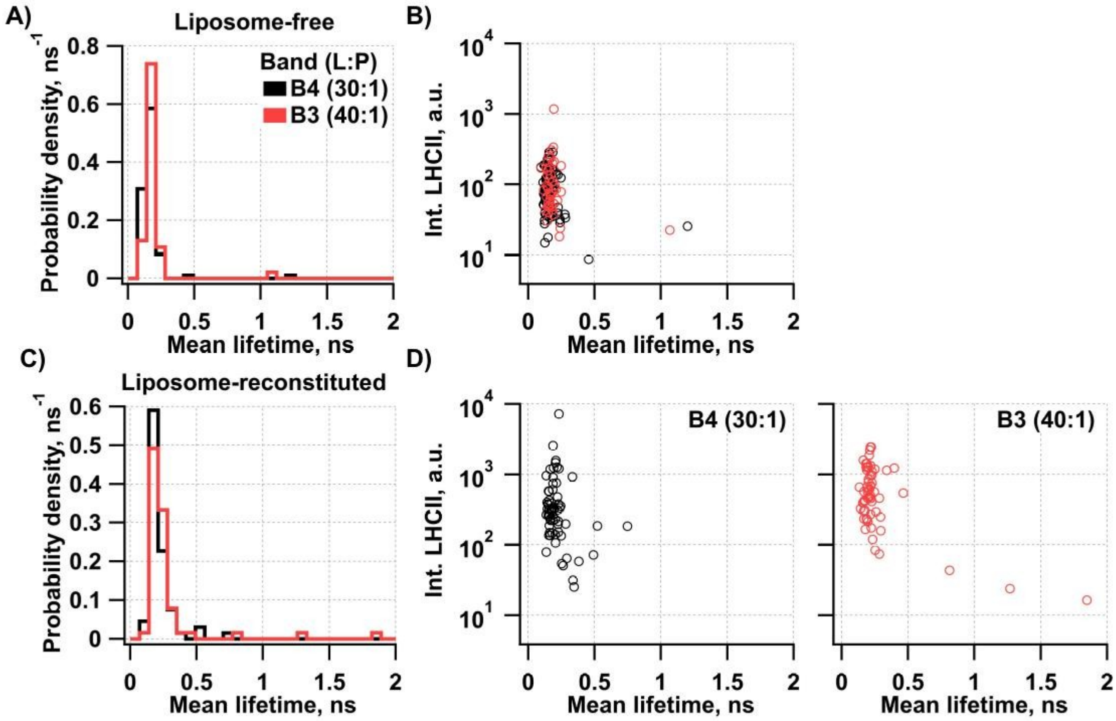


**SI Figure 4** | The results of fluorescence lifetime imaging microcopy. **A)** Graphs showing distribution of fluorescence lifetimes obtained by the single exponential function fitting of liposome-free and **C)** -reconstituted LHCII of two samples with different lipid to protein (L:P) ratios. **B)** Graphs showing correlation of fluorescence lifetimes of liposome-free and **D)** -reconstituted LHCII of three samples with different lipid to protein ratio (L:P).

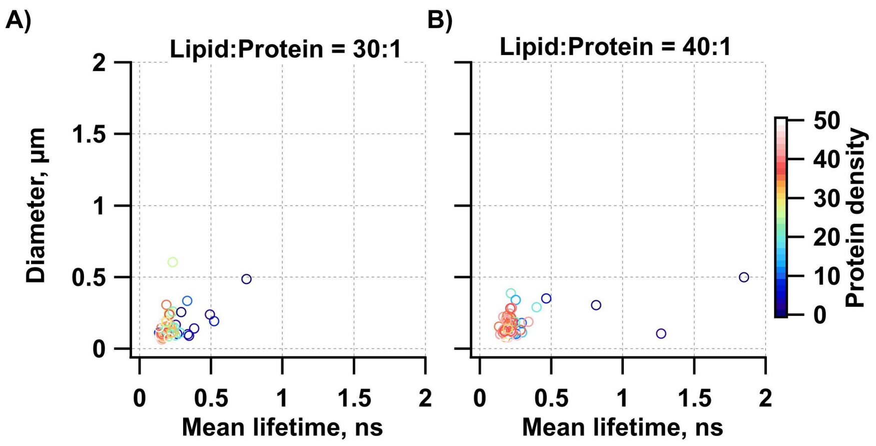


**SI Figure 5** | Correlation of fluorescence lifetimes of liposome-reconstituted LHCII of two samples with different lipid to protein ratio: **A)** 30:1, and **B)** 40:1 that were obtained from 500:1 sample. Color-code represents the protein density.
